# Parallel signatures of mammalian domestication and human industrialization in the gut microbiota

**DOI:** 10.1101/611483

**Authors:** Aspen T. Reese, Katia S. Chadaideh, Caroline E. Diggins, Mark Beckel, Peggy Callahan, Roberta Ryan, Melissa Emery Thompson, Rachel N. Carmody

## Abstract

Domestication may have had convergent effects on the microbiota of domesticates and humans through analogous ecological shifts. Comparing the gut microbiota of domestic and related wild mammals plus humans and chimpanzees, we found consistent shifts in composition in domestic animals and in humans from industrialized but not traditional societies. Reciprocal diet switches in mice and canids demonstrated that diet played a dominant role in shaping the domestic gut microbiota, with stronger responses in the member of the wild-domestic pair with higher dietary and microbial diversity. Laboratory mice recovered wild-like microbial diversity and responsiveness with experimental colonization. We conclude that domestication and industrialization have similarly impacted the gut microbiota, emphasizing the utility of domestic animal models and diets for understanding host-microbial interactions in rapidly changing environments.

---

Changes in industrialized human lifestyles have resulted in large shifts in the gut microbiota relative to traditional populations or closely related primates, including reductions in alpha-diversity and changes in composition (*1-4*) that have been implicated in the rise of various metabolic and immunological diseases (*5-7*). Ecological differences between industrialized humans and chimpanzees, and to a lesser extent between industrialized and non-industrialized human populations, resemble those between domestic and wild animals, including shifts toward non-seasonal calorically-dense diets, reduced physical activity, variations in movement and density, changes in pathogen exposure and antibiotic use, and altered reproductive patterns (*8*). Furthermore, the evolution of *Homo sapiens* has been argued to reflect self-domestication arising due to selection for reduced social aggression (*9*). Despite these parallels, the global effects of domestication on the gut microbiota and its relationship to the effects of human industrialization remain unclear.

Notably, many of the altered ecological features experienced by domesticated animals and industrialized humans have been independently observed to impact the gut microbiota, including diet (*10, 11*) physical activity (*12, 13*), the size and nature of social networks (*14, 15*), antibiotic use (*16, 17*), and changes in birthing and lactation practices (*16, 18*). This overlap leads to the predictions that (i) gut microbial communities will differ between domestic animals and their wild counterparts, (ii) gut microbial communities of diverse domestic animals may exhibit convergent characteristics in a microbial counterpart to the physiological domestication syndrome (*19*), and (iii) gut microbial changes observed with domestication may parallel contrasts observed between chimpanzees and industrialized humans. In addition, to the extent that domestication effects are driven by ecology rather than host genotype, we should expect (iv) humans in traditional and industrialized societies will differ, and (v) experimental control of environmental variables should be able to overcome differences in the gut microbiota between closely related hosts.

Here, we evaluate these predictions by reporting the effects of domestication on the mammalian gut microbiota, comparing these effects to those of human industrialization, and exploring the genetic and ecological forces driving these patterns. First, we characterized the fecal microbiota of wild and domestic populations of nine pairs of artiodactyl, carnivore, lagomorph, and rodent species (Fig. 1A) using 16S rRNA gene amplicon sequencing and qPCR. We found consistent effects of domestication status on gut microbiota composition, despite observing no single convergent profile. Domestication status contributed significantly to variation in microbial communities (P<0.001, R^2^=0.16, PERMANOVA), although the largest single factor was host pair (e.g., pig/boar; P<0.001 R^2^=0.39; Fig. 1B). Diet and digestive physiology were also determinants (P<0.001, R^2^=0.11 diet, R^2^=0.14 physiology; Fig. S1), as seen in other surveys of mammals (*20*), with effect sizes comparable to that of domestication status. Consistent with the idea that higher ecological homogeneity may lead to more similar gut microbial communities in domesticates, we found there was greater between-animal variability in wild gut communities than in domesticates (P=0.005, F=8.833; permutation test for F).

**Fig. 1.**
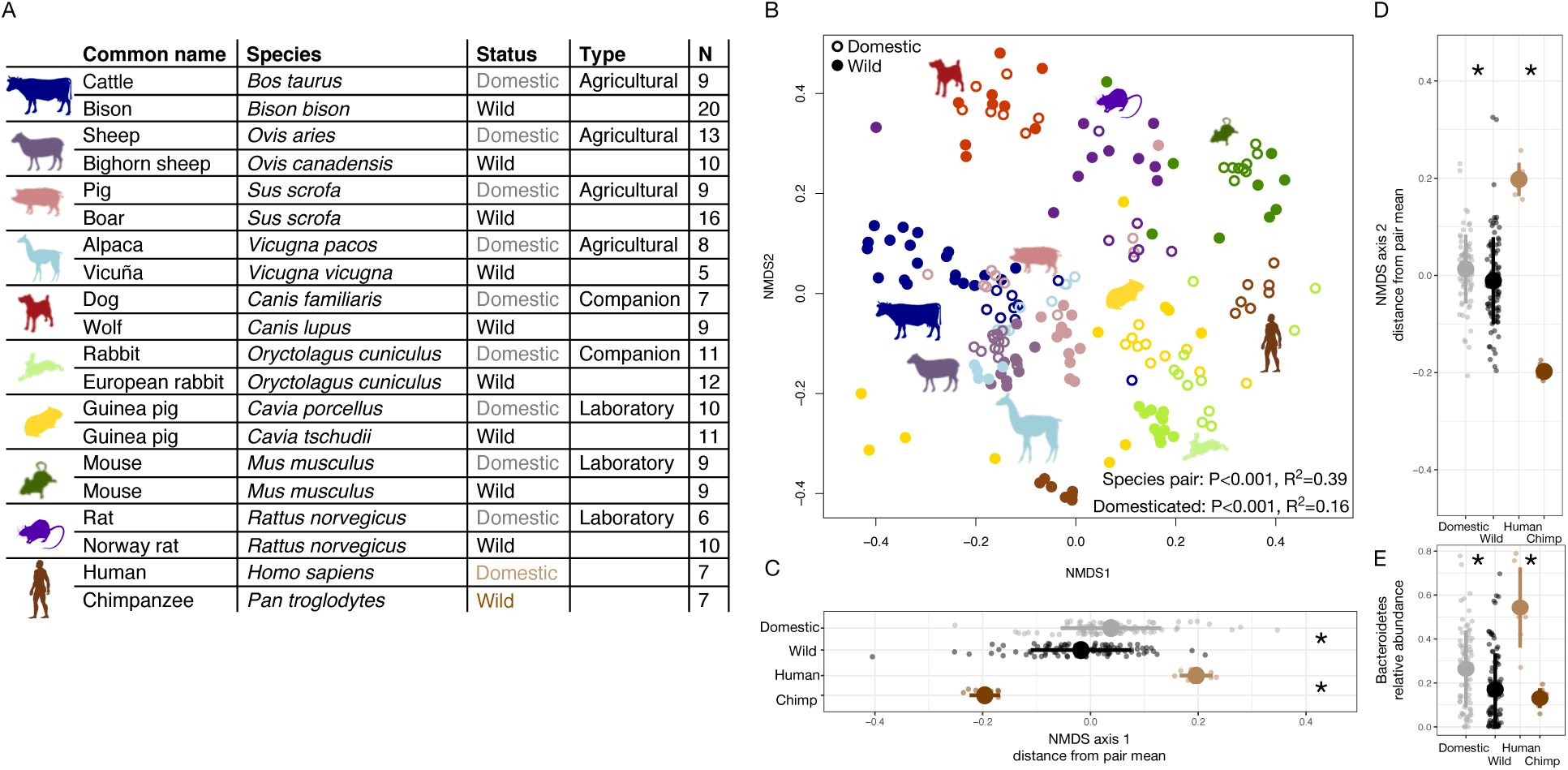
The gut microbiota of wild and domestic mammals differ consistently and in a manner recapitulating differences between industrialized humans and chimpanzees. (**A**) Sampling scheme for cross-species study. (**B**) Nonmetric multidimensional scaling (NMDS) ordination of Bray-Curtis dissimilarities illustrates a significant signal of domestication (closed versus open circles) and clustering by species pair (color). (**C**) Individual shifts relative to species-pair mean along the first NMDS axis differ by domestication status. (**D**) Individual shifts relative to species-pair mean along the second NMDS axis differ by domestication status. (**E**) Relative abundance of the bacterial phylum Bacteroidetes differs by domestication status. Asterisks in (C-E) indicate P<0.05 Mann-Whitney U test by domestication status for animals or by species for human/chimpanzees. Large circles are means; bars show standard deviations.

To determine whether there was a consistent shift in microbial composition with domestication, we calculated the difference between an individual’s ordination coordinates and the average of its host pair along the first and second NMDS axis. Domestic individuals were typically further right (axis 1: P<0.001, Mann-Whitney U test; Fig. 1C) and further up (axis 2: P=0.007; Fig. 1D) relative to the average of their host pair. Domestic species all displayed these shifts, whether classified as laboratory, agricultural, or companion animals (P<0.05, Mann-Whitney U tests; Fig. 1A, S2).

Microbial density quantified as copies of the 16S rRNA gene per gram of feces (P=0.089, Mann-Whitney U test), OTU richness (P=0.800), and Shannon index (P=0.200; Fig. S3) did not differ based on domestication status, indicating that the domestication signal overall is not primarily driven by species loss. By contrast, we observed changes in the abundances of certain microbial taxa. Across host taxa, domestication was associated with higher abundances of the phyla Bacteroidetes (P=0.023, Bonferroni-corrected Mann-Whitney U test; Fig. 1E, S3) and Verrucomicrobia (P=0.001; Fig. S3). These phyla are known to be overrepresented in industrialized compared with traditional human populations (*4*). Consistent with heightened environmental exposure, wild animals generally had more diverse (P=0.001, Mann-Whitney U test) and marginally more abundant (P=0.092; Fig. S3) communities of microbes recognized as potential human pathogens. Among laboratory animals specifically, microbial richness (P=0.045, Mann-Whitney U test), potential pathogen abundance (P<0.001), and pathogen richness (P<0.001) were all substantially lower than among wild relatives, while total microbial load was higher (P=0.006; Fig. S2). Agricultural animals had higher Shannon index values (P=0.001, Mann-Whitney U test) and marginally higher pathogen abundances (P=0.067; Fig. S2) compared with their wild counterparts. By contrast, companion animals did not differ significantly by domestication status for microbial load, diversity, or pathogen metrics. The elevated pathogen abundances found in wild populations overall may largely be ascribed to differences in laboratory animals, which are maintained under conditions that minimize the likelihood of infection. Under natural conditions, however, the domestic microbiota may exhibit reduced colonization resistance or immune system functioning (*21, 22*), resulting in higher pathogen colonization, as observed in agricultural animals.

Given the hypothesis that *Homo sapiens* has undergone a process of self-domestication (*9, 19*), we next tested whether the gut microbial communities of industrialized humans and chimpanzees exhibit parallel shifts to those observed between domestic animals and their wild counterparts when compared in the same ordination space. Indeed, this is what we found (P<0.001, Mann-Whitney U tests; Fig. 1C, 1D). Microbial load (P=0.002, Mann-Whitney U test) and Shannon index (P=0.018; Fig. S3) also differed between industrialized humans and chimpanzees, with industrialized humans harboring microbial communities with substantially lower alpha-diversity. Consistent with the greater evolutionary and profound ecological distance between humans and chimpanzees (*2*), the magnitude of the microbial difference between industrialized humans and chimpanzees exceeded that observed for other animal pairs. To estimate the divergence attributable to ecology versus host genotype, we proceeded to compare the gut microbial communities of humans living in industrialized versus traditional societies. Reanalysis of our cross-species comparison to include published data on human populations in rural Malawi and Venezuela (*23*) (see *Methods*) found that the gut microbial communities of these traditional populations differed substantially from those of two independent U.S. samples, clustering more closely to those of chimpanzees (Fig. S4). These data indicate that the human gut microbiota does not carry a global signal of domestication, as would be predicted under the human self-domestication hypothesis. Rather, they suggest that gut microbial responses to domestication and industrialization are more likely driven by common ecological factors, a conclusion further supported by the observation that domestic animals were significantly more similar to those of industrialized humans than their wild animal counterparts (P=0.002, Mann-Whitney U test). Notably, the gut microbial communities of domestic animals and industrialized humans most closely resembled one another for companion and laboratory animals (P<0.001, Kruskal-Wallis test; Fig. S2), presumably reflecting their greater degree of overlap in ecological variables and physical contact (*24*).

Importantly, the observation that gut microbial divergence is restricted to industrialized populations implicates recent ecological changes as opposed to ecological changes with deeper roots in human evolution. Many recent ecological changes involve accelerations of basic patterns established during the evolution of *Homo*, including increased proportion of calories from fat and protein, increased dependence on animal source foods, and extensive food processing by thermal and non-thermal means (*25*). Other ecological changes are likely specific to industrialization, including reduced physical activity and antibiotic use. Further work will be required to illuminate the combination of ecological factors driving similarities between the domesticated and industrialized microbial profiles.

To begin to tease apart these ecological drivers, we performed a series of reciprocal diet experiments that tested the extent to which gut microbial signatures of domestic-wild pairs could be recapitulated and reversed solely by the administration of domestic versus wild diets. We first conducted a fully factorial experiment in which wild-caught and laboratory mice (*Mus musculus*) were maintained for 28 days on wild or domestic diets (Fig. 2A, Table S1). Overall, we found that host genotype explained the largest amount of variation in composition (P<0.001, R^2^=0.173, PERMANOVA), but diet (P<0.001, R^2^=0.042) and a genotype by diet interaction term (P<0.001, R^2^=0.020) were also significant (Fig. 2B, S5). Experimental groups varied in their microbial responsiveness over the course of the experiment (axis 1: P=0.063, axis 2: P<0.001, Kruskal-Wallis tests; Fig. 2C, S5). Generally, the microbiota of Wild_G_/Dom_D_ mice moved toward the Dom_G_/Dom_D_ mouse average community, the Dom_G_/Wild_D_ microbiota moved in the opposite direction, and those of Wild_G_/Wild_D_ and Dom_G_/Dom_D_ mice did not shift (Fig. 2B). Over the course of the experiment, Shannon index values also changed significantly across treatment groups (P=0.005, Kruskal-Wallis test), with Dom_G_/Wild_D_ mice becoming significantly more diverse (P=0.002, one-sample Wilcoxon test) despite initial differences in alpha-diversity between wild and domestic mice (P=0.009, Mann Whitney U test; Fig. S6).

**Fig. 2.**
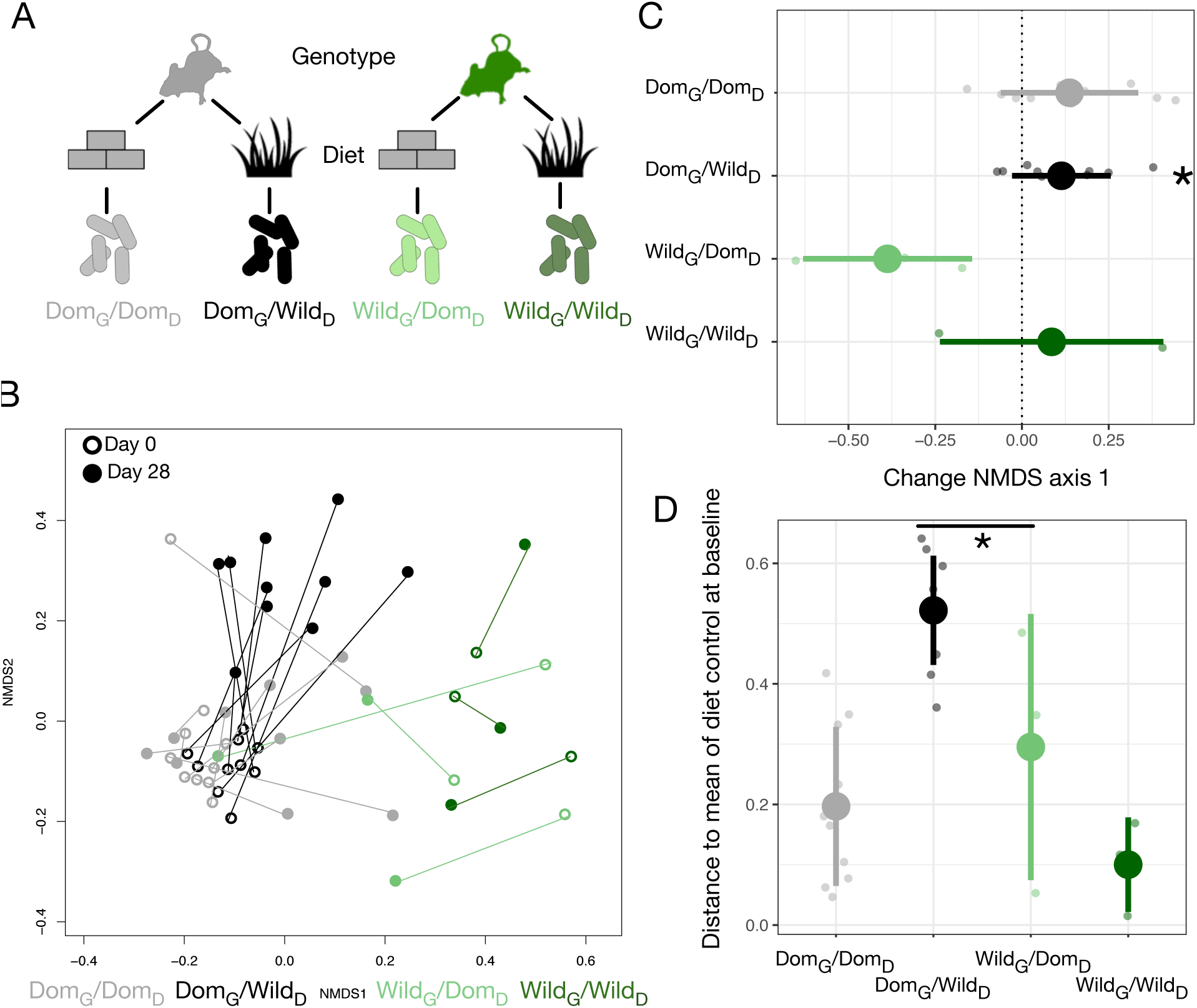
Microbial differences between wild and domestic mice can be overcome by diet shifts. (**A**) Design scheme for genotype/diet factorial mouse experiment. (**B**) Nonmetric multidimensional scaling (NMDS) ordination of Bray-Curtis dissimilarities showing changes for mice from day 0 (open circle) to day 28 (closed circle) by experimental groups (color). (**C**) Animals on reciprocal diets (Dom_G_/Wild_D_ and Wild_G_/Dom_D_) move in opposite directions along NMDS axis 1 from day 0 to day 28. Asterisk indicates P<0.05 one-sample Wilcoxon test. Dashed line indicates a shift of 0. (**D**) At the end of the experiment, distance to the mean of the diet control at baseline (Dom_G_/Dom_D_ and Wild_G_/Wild_D_) was lower for wild mice than lab mice. Asterisk indicates P<0.05 Mann-Whitney U test. Large circles are means; bars show standard deviations.

Neither diet nor host genotype were associated with differences in microbial density over the experiment (P=0.272, Kruskal-Wallis test; Fig. S6), but it is notable that the total amount of feces produced, and thus likely the total number of bacteria, was lower in each host genotype when fed wild diet (P<0.001, Kruskal-Wallis test; Fig. S6). Despite similar trends in fecal production between the experimental groups, energy harvest responses differed markedly between experimental groups (P<0.001, Kruskal-Wallis test; Fig. S6). While wild mice were equally efficient consumers of both diets, laboratory mice captured 15% fewer calories when consuming the wild versus domestic diet. Nonetheless, weight gain in laboratory mice did not differ between diet groups, while Wild_G_/Dom_D_ mice tended to gain weight over the course of the experiment (P=0.250, one-sample Wilcoxon test; Fig. S6). Interestingly, the asymmetry in energy harvest between genotypes was also reflected in differential microbial responses to reciprocal diets. Whereas the microbial communities of Wild_G_/Dom_D_ mice eventually largely recapitulated those of untreated Dom_G_ mice, the microbial communities of Dom_G_/Wild_D_ mice remained distinct from untreated Wild_G_ mice throughout the experiment (P=0.042, Mann-Whitney U test; Fig. 2B). The inability to foster a wild-type microbiota may underpin the reduced digestive efficiency of the Dom_G_/Wild_D_ mice.

We hypothesized that these asymmetries were due to past extinction of relevant strains from laboratory microbial communities and no dispersal source of replacement strains (*26*). Therefore, we tested whether experimental dispersal from a wild microbial community in conjunction with feeding a wild diet could support a fully wild microbial community in laboratory mice (Fig. 3A). A single colonization treatment with a wild mouse cecal community (via gavage) led to significant shifts in the microbial community (Fig. 3B, S7), resulting in closer resemblance to the wild donor (P<0.001, Mann-Whitney U test; Fig. 3C). While laboratory mice fed a wild diet but given a control gavage (PBS) also moved toward the donor along NMDS axis 1 (P=0.002, one-sample Wilcoxon test; Fig. 3D), reflecting the influence of diet, the magnitude of the shift following the experimental colonization was substantially greater (P<0.001, Kruskal-Wallis test). There were no apparent differences in these shifts based on diet treatment among colonized mice (P=0.182, Mann-Whitney U test). Colonization with a wild community led to an increase in alpha-diversity as measured by the Shannon index (P=0.042, Kruskal-Wallis test), and wild diet treatment led to reductions in fecal production (P<0.001; Fig. S7). Although all mice exhibited an increase in load over the course of the experiment (P<0.05, one-sample Wilcoxon tests), colonization with a wild community did not lead to higher loads overall (P=0.742, Kruskal-Wallis test; Fig. S7). This result suggests that differences observed with treatment reflected shifts in gut microbial community structure rather than simple augmentation.

**Fig. 3.**
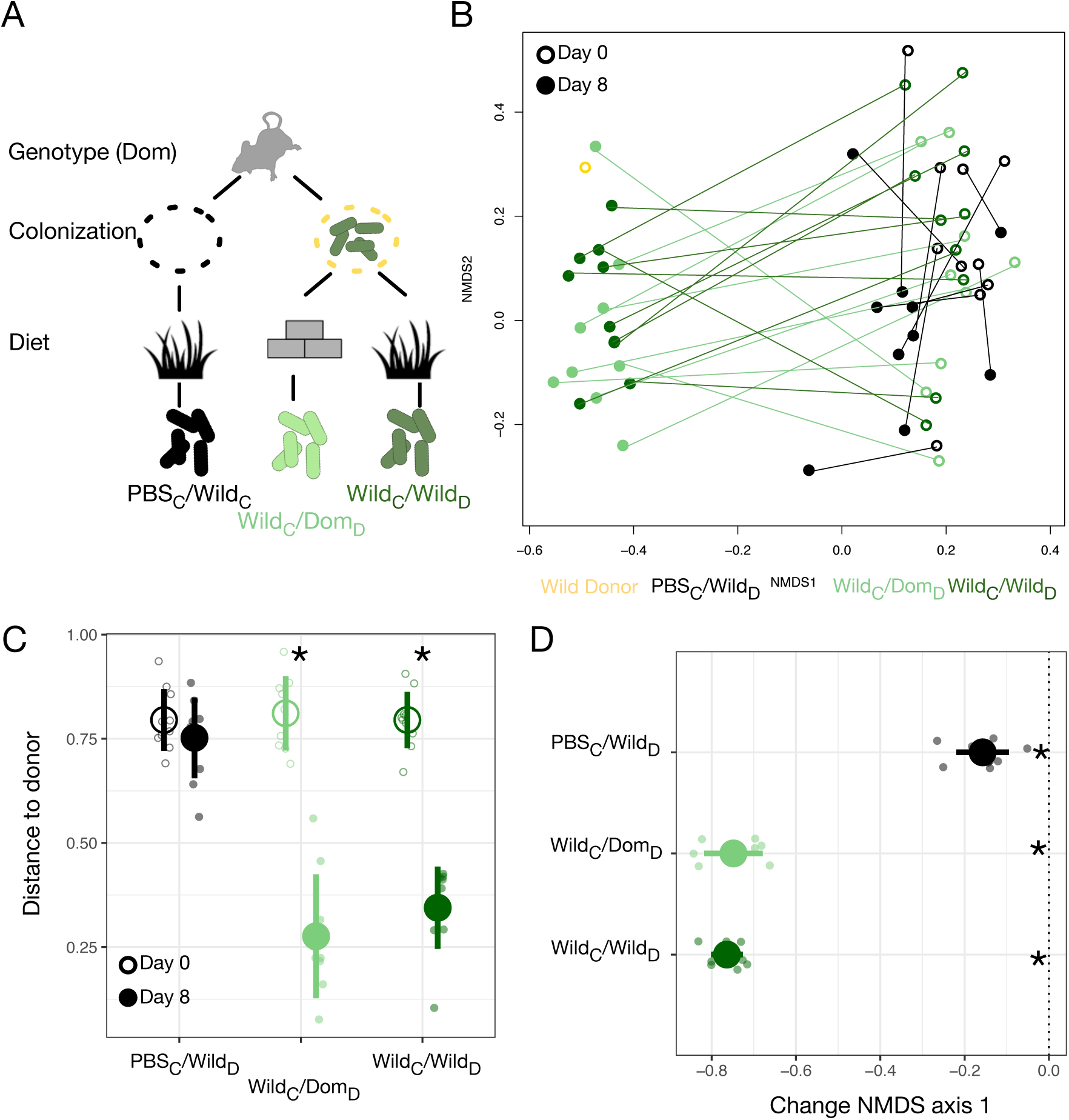
Laboratory mice can be re-wilded through colonization with wild microbial community. (**A**) Design scheme for colonization/diet mouse experiment. (**B**) Nonmetric multidimensional scaling (NMDS) ordination of Bray-Curtis dissimilarities showing changes for mice from day 0 (open circles) to day 8 (closed circles) by experimental groups (color). (**C**-**D**) At the end of the experiment (closed circle), distance to the wild community donor decreased most in animals colonized with wild communities (Wild_C_/Dom_D_ and Wild_C_/Wild_D_; C), but all experimental groups exhibited change along NMDS axis 1 (D) during the course of the experiment. Asterisks in (C) indicate P<0.05 Mann-Whitney U test comparing day 0 to day 8 for each experimental group. Asterisks in (D) indicate P<0.05 one-sample Wilcoxon test, and dashed line indicates a shift of 0. Large circles are means; bars show standard deviations.

To test if these findings were generalizable to non-laboratory animals, we conducted an analogous reciprocal diet experiment in a captive sympatric population of wolves and dogs (Fig. 4A). We tracked gut microbial dynamics in these canids for one week on their standard diet (raw carcasses or commercial dog food, respectively) and one week on the reciprocal diet. As in the mouse experiment, we found that host genotype explained the largest amount of variation in gut microbiota composition (P<0.001, R^2^=0.098, PERMANOVA), but diet (P<0.001, R^2^=0.058) and a genotype by diet interaction term (P<0.001, R^2^=0.028) were also significant (Fig. 4B, S8). There were significant differences between experimental groups in the magnitude of their shifts along the first (P<0.001, Kruskal-Wallis test; Fig. 4C) and second (P=0.045; Fig. S8) NMDS axes over the experimental periods. As in the mouse experiments, we observed animals on reciprocal diet treatments moved significantly toward the diet control of the other species (P<0.05, one-sample Wilcoxon tests; Fig. 4B), while the control animals did not shift predictably (P>0.100).

**Fig. 4.**
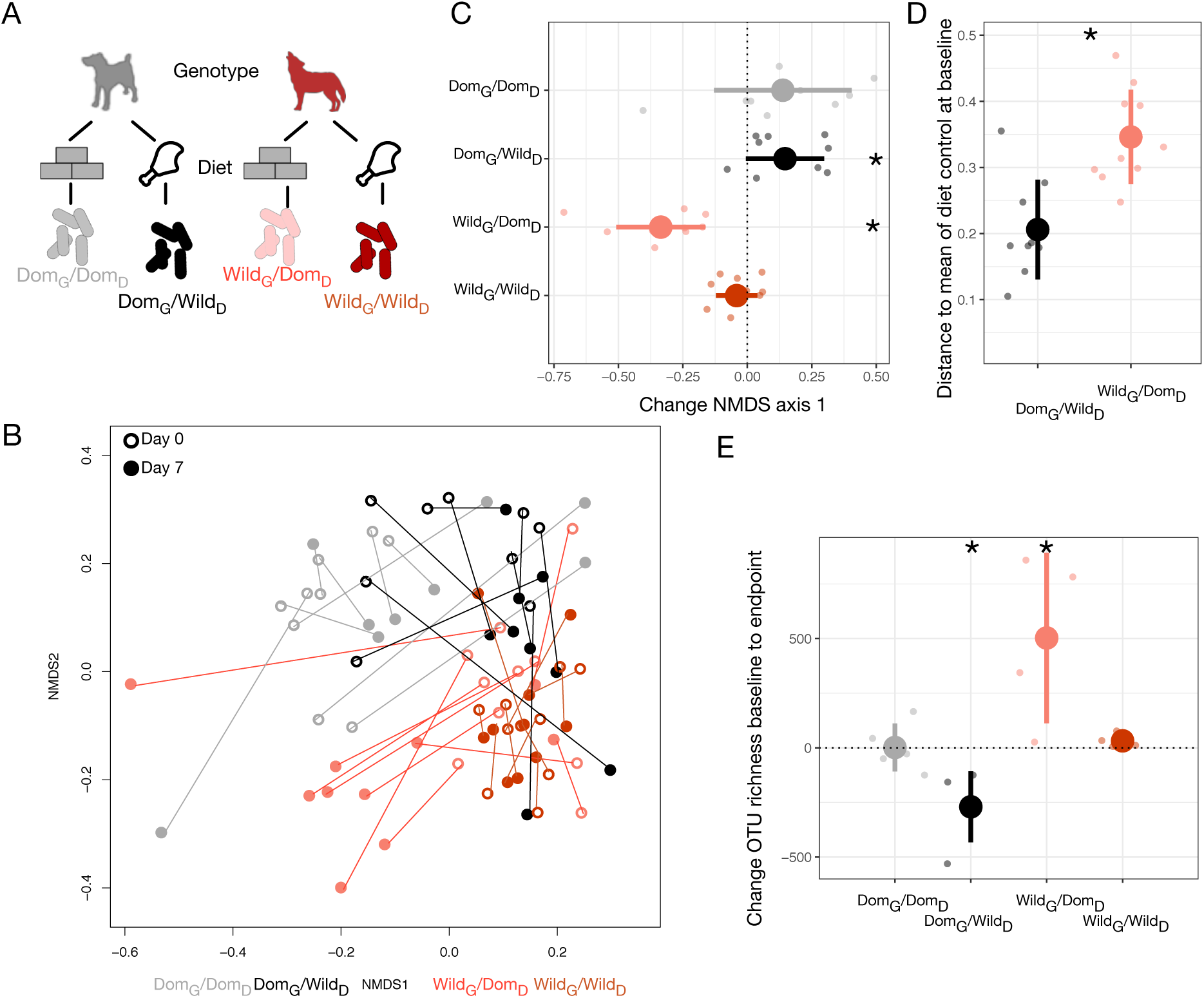
Microbial differences between wild and domestic canids were overcome by diet shifts, as in mice. (**A**) Design scheme for genotype/diet canid experiment. (**B**) Nonmetric multidimensional scaling (NMDS) ordination of Bray-Curtis dissimilarities showing changes for canids from day 0 (open circle) to day 7 (closed circle) by experimental groups (color). (**C**) Canids on reciprocal diets (Dom_G_/Wild_D_ and Wild_G_/Dom_D_) moved in opposite directions along NMDS axis 1 over time. (**D**) At the end of the experiment, distance to the mean of diet controls at baseline (Dom_G_/Dom_D_ and Wild_G_/Wild_D_) was lower for dogs than wolves on reciprocal diets. Asterisk indicates P<0.05 Mann-Whitney U test. (**E**) Change in OTU richness from day 0 to day 7 differed significantly from 0 in opposite directions for animals on reciprocal diets (Dom_G_/Wild_D_ and Wild_G_/Dom_D_). Asterisks in (C, E) indicate P<0.05 one-sample Wilcoxon test, and dashed line indicates a shift of 0. Large circles are means; bars show standard deviations.

Again, we observed an asymmetry in the degree of microbial composition change between domestic and wild animals. On experimental diets, dogs and wolves differed significantly in their dissimilarity to diet controls (P<0.001, Kruskal-Wallis test, Fig. 4D), with the gut microbial communities of dogs fed raw carcasses resembling those of wolves at baseline but the gut microbial communities of wolves fed dog food remaining distinct from those of dogs at baseline (P=0.001, Mann-Whitney U test). The difference in the direction of asymmetry between the mouse and canid experiments may be explained by the different trends in the diet ecology between omnivores and carnivores during domestication. Carnivores, through the addition of extensive carbohydrates to their diet (*27*), likely encounter more diverse diets in captivity than in the wild, whereas herbivores and omnivores eat a smaller number of plant species or even just a single feed mix. Supporting this, we found dogs initially had significantly higher OTU richness (P<0.001) and Shannon index (P=0.003) than wolves (Fig. S9), but that reciprocal diets led to a switch in diversity (richness: P=0.014, Shannon index: P=0.027, Mann-Whitney U tests), with wolves becoming more diverse on dog food while dogs lost diversity on raw carcasses (Fig. 4E).

Our reciprocal diet experiments in mice and canids confirm that ecology plays a predominant role in shaping the domestic gut microbiota. Moreover, that the effects of a single ecological variable like diet were sufficiently profound to outweigh those of host genotype suggests that suites of ecological variables changing together, such as during domestication or industrialization, may have collectively exerted an even larger influence. However, microbiota changes were certainly not the only pathway for domesticating animals to respond to changing ecological factors. For example, in dogs, genetic changes have enhanced starch digestion (*27*). The increased microbial diversity and shifts in microbial composition that we observed in dogs may likewise contribute to carbohydrate digestion and may have been particularly important early in domestication, before host evolution occurred, although that hypothesis remains to be tested. Notably, the microbiota has been found to supplement evolutionary responses during dietary niche expansion in wild animals that consume plants high in toxins (*28*). As such, the changes observed in domestic animals are not necessarily maladaptive, as the industrialized human microbiome is often characterized to be (*29*). Beyond host support of microbiota that can better digest a domestic diet, humans may have selected for animals harboring a microbiota that helped them grow and reproduce well on such diets. Specialization for microbial performance domestic diets may have come at the cost of broader digestive capacity, as seen in the domestic mouse microbiota, which was better at harvesting energy from domestic diets than from wild diets (Fig. S6). Future studies examining the trade-offs between microbially-mediated functions, like digestive capacity, reproduction, and immunity, will help to illuminate the complex selection pressures shaping the domestic holobiont.

Taken together, our data reveal strong parallels between the gut microbial signatures of domestication and industrialization, most likely driven by convergent changes in ecology, including diet. Because laboratory mice demonstrate some of the largest overall differences relative to their wild counterparts, and in part emulate the variation observed between industrialized humans and closely related primates, their translational potential as models for studying the gut microbiota of industrialized populations may be greater than currently appreciated. However, our data also suggest that laboratory animals may not be broadly representative of natural host-microbe interactions or their evolutionary history (*30*). Nevertheless, that laboratory mice were permissive of recolonization by wild strains indicates that the local extinctions that occurred during domestication and/or generations in captivity can potentially be mitigated. Previous work has relied on germfree mice colonized with a wild microbiota but fed standard laboratory chow (*21*). A combination of these approaches— adding wild community members and feeding wild diet—would be expected to best support a wild microbiota in laboratory mice. A wild-microbiota laboratory-genotype model could be especially useful for studying infection challenges, disentangling host gene versus microbiota contributions to disease phenotypes, and testing for coevolution between host and microbes.

More generally, our data add to growing evidence that the gut microbiota is finely tuned to variations in the environment, affording at once an opportunity for host-microbial mismatch and an opportunity for rapid microbiota-mediated host adaptation to novel environments (*31*). Further work to characterize the ecological significance of gut microbial plasticity will help reveal the fundamental nature of the host-microbial relationship, the conditions under which plasticity is beneficial versus detrimental, and the ecological conditions promoting cooperative, commensal, and competitive dynamics.

## Materials and Methods

### Fecal sample collection

Gut microbiota samples from a range of non-human species were collected by authors or collaborators primarily from feces. Fecal samples from non-human mammals were collected from the ground within seconds to hours of production. In the case of artiodactyl, carnivore, lagomorph, and rodent feces, this approach precluded the need for institutional approval. Chimpanzee fecal samples were collected under the approval of the UNM IACUC (Protocol 18-200739-MC) and with permission of the Uganda Wildlife Authority and Uganda National Council for Science and Technology. Human samples were self-collected by healthy study participants after providing written informed consent under the approval of the Harvard University IRB (Protocol 17-1016). All samples were flash-frozen or preserved in ethanol prior to permanent storage at −80°C.

#### Domestic animals

Domestic sheep (*Ovis aries*; N=11, 10 female), cattle (*Bos taurus*; N=10, sex unknown), and pig (*Sus scrofa domesticus*; N=9, sex unknown) fecal samples were collected from a farm in Vershire, Vermont. Domestic alpaca (*Vicugna pacos*; N=8, sex unknown) and domestic sheep (*Ovis aries*; N=2, 2 female), fecal samples were collected from a farm in Groton, Massachusetts. Domestic rabbit (*Oryctolagus cuniculus*; N=11, 4 female) fecal samples were collected from a shelter in Billerica, Massachusetts. Mouse (*Mus musculus*, N=9, 0 female), rat (*Rattus norvegicus*; N=6, sex unknown), and guinea pig (*Cavia porcellus;* N=10, 0 female) fecal samples were collected from animals in Harvard laboratory facilities. Dog (*Canis lupus familiaris*; N=7, 4 female) fecal samples were collected from personal pets in Stacy, Minnesota.

#### Wild animals

Wild boar (*Sus scrofa*; N=16, 5 female) fecal samples were collected from adults and juveniles in southeastern Alabama during fall 2017. Rat (*Rattus norvegicus*; N=10, 3 female) gut samples from adults and juveniles were collected directly from the colon shortly following trapping in New York City between February and May 2017 (*32*). Bison (*Bison bison*, N=20, sex unknown) fecal samples were collected from a semi-free-ranging population in Elk Island National Park, Alberta, Canada (*33*). Wild house mouse (*Mus musculus*, N=9, sex unknown) fecal samples were collected from live-trapped animals in the Boston, Massachusetts area during winter 2018. Pursuant to Massachusetts state law, permits were not necessary to trap animals indoors. Wild European rabbit (*Oryctolagus cuniculus*; N=12, sex unknown) fecal samples were collected in Mértola, Portugal during spring 2018. Bighorn sheep (*Ovis canadensis*; N=10, sex unknown) fecal samples were collected during 2017 and 2018 in Wyoming. Vicuña (*Vicugna vicugna*; N=4, 2 female) fecal samples were collected during spring 2018 from a captive population in Santiago, Chile that was free-grazing but supplemented with hay. Wild guinea pig (*Cavia tschudii*, N=11, sex unknown) fecal samples were collected at a facility in Lima, Peru during spring 2018. Wolf (*Canis lupus*; N=9, sex unknown) fecal samples were collected during fall 2017 from captive packs at the Wildlife Science Center in Stacy, Minnesota fed an exclusively raw diet. Wild chimpanzee (*Pan troglodytes schweinfurthii*, N=7, 7 female) fecal samples were collected between September 2015 and January 2016 from adult members of the Kanyawara community in Kibale, Uganda.

#### Human

Fecal samples were collected from healthy adult humans (N=7, 5 female) residing in the Cambridge, Massachusetts area. All participants were provided with sterile study kits, and self-collected fecal samples during the same 3-day period in December 2017. During this period, participants freely consumed their habitual diets. Fecal samples were immediately stored at −20°C and were transferred within 24 hours to permanent storage at −80°C.

#### Human sample meta-analysis

To compare the microbial differences between wild and domestic animals or US humans and chimpanzees with other human populations, we also performed analyses including all of the samples outlined above and a subset of published data from Yatsunenko and colleagues (*23*). We subsampled 7 adult females from their Malawian, Venezuelan, and American populations, downloading the data from MG-RAST. All sequences were trimmed to 100 bp before analysis (see 16S rRNA gene analysis below), and the published dataset was rarefied to 100,000 reads per sample to ensure comparable sequencing depth with our data.

### Animal experiments

#### Wild mouse capture

*Mus musculus* were introduced to North America from Western Europe and are now commonly found in commensal settings (*34*). We set out Sherman live traps in the evenings in buildings and barns during February 2018. Traps were baited with peanut butter and a chunk of fruit and outfitted with sufficient bedding and food to sustain an adult mouse for at least 48 hr. They were checked the following morning to minimize time spent in the traps. Rodents were immediately transferred from their traps to a plastic bag, and unwanted rodent species were released immediately. Mice that were identified as *Mus musculus* (rather than *Peromyscus spp*., also common in Massachusetts) were transferred to temporary cages for transport to lab facilities. At time of capture, we collected fecal samples and body swabs for zoonoses testing by Charles River. The only agent of concern found was fur mites. Because animals were not treated for parasites or pathogens in order to increase maintenance of the wild-state microbiota, they were housed under non-SPF conditions at Harvard’s Concord Field Station. Mice were allowed at least three days to adjust to laboratory conditions without handling and provided with a wild mouse diet [a mix of bird seed (Wagner’s Eastern Regional Blend Deluxe Wild Bird Food) and freeze-dried mealworms; Table S1] before the beginning of the experiment. All mice were housed singly from the time of arrival at the Concord Field Station and had access to water and food ad libitum.

#### Wild/laboratory mice reciprocal diet experiment

A total of 10 wild mice were captured for this experiment. Of these, 2 were deemed too young for inclusion in the study, 1 died before beginning the experiment, and 1 died during the course of the experiment. As a result, we collected 6 wild mice (Wild_G_) that were included in the full study. In addition to the wild mice, male C57BL/6 mice 10-12 weeks of age with a conventional microbiota were purchased from Charles River Laboratories for inclusion in the study (Dom_G_). All mouse experiments were conducted in accordance with the National Institutes of Health Guide for the Care and Use of Laboratory Animals using protocols approved by the Harvard University Institutional Animal Care & Use Committee (protocol number 17-11-315). All mice were housed singly from the time of arrival at the Concord Field Station and had access to water and food ad libitum. Mice were provided nesting material and plastic enrichment housing atop corncob bedding. The mice were maintained in a room with natural light cycles kept at 20-22°C.

Mice, both wild and laboratory, were randomly assigned to one of two dietary treatment groups (N=10 laboratory mice or 3 wild mice per group). The first group (domestic diet: Dom_D_) was provided *ad libitum* mouse chow (Prolab Isopro RMH 3000) in hanging food hoppers, as is standard in mouse studies. The second group (wild diet: Wild_D_) was provided a mix of bird seed (Wagner’s Eastern Regional Blend Deluxe Wild Bird Food) and freeze-dried mealworms (Table S1) in excess of predicted consumption. The food was placed in the corncob bedding to simulate foraging.

Before initiating the dietary interventions, all individuals were weighed and multiple fecal samples were collected. The mice were then returned to a new, clean cage with the treatment diet present. Over the next week, fecal samples and weights were collected daily for each mouse. The amount of food remaining was weighed and additional wild diet was added daily. One week after beginning the experiment, mice were weighed and fecal samples collected then mice were moved to clean cages. Weights and fecal samples were henceforth collected weekly (day 14, 21, 28) until the end of the experiment, although additional food was added biweekly for individuals assigned to the wild diet treatment. Additional chow was added to hoppers for individuals assigned to the conventional diet treatment, and all water bottles were refilled as necessary. At the end of each week, cage bedding was collected and sifted to quantify uneaten food (Wild_D_) and total weekly fecal production (all groups during week 3), as well as to provide fecal samples for bomb calorimetry (6050 Calorimeter, Parr). All calorimetry results were adjusted for the average weekly fecal production and average weekly food intake of each experimental group. At the end of the experiment (day 28-30), mice were humanely sacrificed via CO_2_ euthanasia.

#### Wild/laboratory mice gavage experiment

Thirty 10 week old male C57BL/6 mice with a native microbiota were purchased from Charles River Laboratories for inclusion in the study. Mice were cohoused in litter groups of 3-4 until beginning the study. Cage groups were spread across the treatment groups, with individuals randomly assigned to a diet and colonization treatment. There were three treatment groups: wild colonized/wild diet (Wild_C_/Wild_D_); wild colonized/domestic diet (Wild_C_/Dom_D_); or PBS gavage/wild diet (PBS_C_/Wild_D_). The latter served as a colonization control, emulating the Dom_G_/Wild_D_ group from the reciprocal diet mouse experiment.

On the first day of study, fecal samples were collected from each mouse and the mice were weighed before colonization. For mice receiving a wild microbiota, we experimentally colonized them with cecal contents collected from one randomly selected Wild_G_/Wild_D_ individual in the wild/laboratory experiment (see above). The cecal contents were prepared following (*21*). In short, frozen cecal contents were resuspended in reduced PBS (1:1 g:ml) under anaerobic conditions then diluted 1:30. Each recipient mouse received a single dose of 100 to 150ul cecal solution via oral gavage. PBS control mice received 100 to 150ul reduced PBS via oral gavage.

Following gavage, mice were transferred to single housing in new, clean cages with the treatment diet present. Mice receiving domestic diet were provided ad libitum mouse chow (Prolab Isopro RMH 3000) in hanging food hoppers. Wild mouse diet consisted of a mix of bird seed (Wagner’s Eastern Regional Blend Deluxe Wild Bird Food) and freeze-dried mealworms (Table S1), which was provided in excess of predicted consumption and placed in the corncob bedding to simulate foraging. All mice were provided with nesting material and plastic enrichment housing atop corncob bedding.

Additional fecal samples and weights were collected on days 1, 2, and 8 following gavage. After weights and fecal samples were collected on day 8, mice were humanely sacrificed via CO_2_ euthanasia. At the end of the experiment, cage material was collected and sifted to quantify uneaten food (Wild_D_) and total weekly fecal production (all groups).

#### Wolf/dog reciprocal diet experiment

Ten wolves (*Canis lupus*) and nine dogs (*Canis familiaris*) participated in the study. Wild-caught or captive born wolves lived in packs ranging in size from 2-6 at the Wildlife Science Center (WSC; Stacy, MN). They were exposed to natural light cycles and weather conditions, with access to shelters and wolf-dug dens in their enclosures. Wolves had ad libitum access to water. Dogs enrolled in this study were privately owned and were recruited to participate through their owners. Dogs were kept in their typical environment throughout the experiment. All canid experimentation was approved by the WSC IACUC (protocol number HAR-001). Wolves were enrolled in the study from Dec. 5 – Dec. 20 2018; dogs from Dec. 24 2018 – Jan. 8 2019.

Every day of the study, animals were given inert glass beads via treats (∼15g raw meatballs for wolves). The beads can be passed naturally without harm to the animal and allowed for source identification for fecal samples in cohoused animals. Fecal samples were collected daily in a sterile manner then moved to −20°C storage before long-term storage at −80°C. For the first week of the experiment all animals received a control diet that matched their genetic background (Table S1)— raw chicken parts (4lbs/animal) for wolves (Wild_G_/Wild_D_) and commercial dog food (Nutrisource Lamb Meal and Peas Grain Free) for dogs (Dom_G_/Dom_D_). Fecal samples were collected at least once daily from wolf enclosures and the dogs’ home environments without handling the animals. On day 8, wolves were provided no new food, but were able to complete consumption of previously provided diet materials. Fecal samples collected on this day were considered baseline samples for the next arm of the experiment. Beginning on day 8, a week of reciprocal diet feeding was commenced. During this period, wolves were fed commercial dog food (Wild_G_/Dom_D_) and dogs were fed raw chicken parts (Dom_G_/Wild_D_); glass beads continued to be administered via treats thus wolves received small amounts (∼15g) of raw meat daily. Daily fecal samples were again collected. Following completion of the study, animals were returned to their standard diet.

### 16S rRNA gene analysis

#### Extraction

Following collection during observational or experimental animal work, fecal samples were temporarily stored at −20°C then moved to −80°C for long term storage. Individual mouse pellets or approximately 0.1g feces were used for DNA extraction using the E.Z.N.A. Soil DNA Kit (Omega) following manufacturer’s instructions.

#### Sequencing

We performed 16S rRNA gene amplicon sequencing on fecal samples to determine gut microbial community structure. We used custom barcoded primers (*35*) targeting the 515F to 806Rb region of the 16S rRNA gene following published protocols (*35-37*). Sequencing was conducted on an Illumina HiSeq with single end 150bp reads in the Bauer Core Facility at Harvard University. Data was processed using Qiime1.8 commands for closed reference OTU picking with 97% similarity. Microbial taxonomy was assigned in reference to the GreenGenes database. We obtained 158611±109567 assigned reads per sample.

#### qPCR

To estimate total bacterial load, quantitative PCR (qPCR) was performed on fecal DNA using the same primers as used for sequencing. qPCR assays were run using PerfeCTa SYBR Green SuperMix Reaction Mix (QuantaBio) on a BioRad CFX384 Touch (Applied Biosystems, Foster City, CA) in the Bauer Core Facility at Harvard University. Cycle-threshold values were standardized against a dilution curve of known concentration and then adjusted for the weight of fecal matter extracted.

### Statistical analyses

All statistical analyses were carried out in R (R core team, ver. 3.3). Alpha-diversity (Shannon index, OTU richness) and beta-diversity (Bray-Curtis) metrics were calculated using the vegan package (*38*). All statistical tests performed were non-parametric. Permutational MANOVA (PERMANOVA) was carried out with the adonis function in vegan. Variability in a species’ microbial community composition was calculated with the permutest and betadisper functions in vegan. For changes in phylum-level abundance, relative abundance data were multiplied by bacterial load measurements; a Bonferroni correction for multiple hypothesis correction was then applied to all test results. Phyla were included if they had an average abundance of at least 0.01% across all samples.

Potential human pathogens were identified following published methods (*39, 40*). In short, we obtained a list of potential human pathogens, compiled by Kembel and colleagues (*39*), then manually compared that list to the taxa identified to genus or species level in analysis. A subset of the data containing only these species was then analyzed for diversity with the same methods used for the total dataset.

To determine the consistency of gut microbial shifts with domestication or industrialization in the observational study, we calculated the average of the species pair (e.g., pig/boar) for axis 1 and axis 2 of the NMDS then measured the shift along each axis for an individual sample and tested for differences by domestication status. To estimate the direction and magnitude of changes in beta-diversity during the experimental studies, we calculated the distance along axis 1 or 2 of the NMDS relative to a baseline sample for that individual. We estimated the direction and magnitude of dissimilarity from the expected community composition (donor microbial community in gavage experiment; baseline species average for Dom_G_/Dom_D_ or Wild_G_/Wild_D_ in diet experiments) as the length of the vector through the first two axes of ordination space.

## Acknowledgments

We thank many collaborators for help in collecting wild and domestic animal fecal samples, including Gwynne Durham and the Mountain School (cattle, pigs, sheep); Luina Greine Farm (alpaca); Steve Ditchkoff (wild boar); Jason Munshi-South and Matthew Combs (rats); Pedro Monterroso, Marisa Rodrigues, and Marketa Zimova (European rabbits); Margaret Gruen and Kyle Smith (dogs, wolves); Kevin Monteith (Bighorn sheep); J. Scott Weese (bison); Cristián Bonacic (vicuña); Bridget Alex, Nicholas Holowka, Irene Li, Daniel Lieberman, Mark Omura, and Antonia Prescott (wild mice). For help in collecting wild chimpanzee samples we thank Richard Wrangham, Zarin Machanda, Martin Muller, Emily Otali, and the staff of Kibale Chimpanzee Project. For assistance in carrying out experiments we thank Cary Allen-Blevins, Rachel Berg, Andy Biewener, Meg Callahan-Beckel, Brian Hare, Kathleen Pritchett-Corning, Pedro Ramirez, Laura Schell, and Emily Venable. For sequencing assistance we thank Christian Daly, Claire Reardon, and The Bauer Core Facility. For helpful comments on the manuscript we thank Daniel Lieberman, Richard Wrangham, and members of the Carmody lab.

## Author contributions

A.T.R. designed the study, conducted analyses, and wrote the manuscript; K.S.C., C.E.D., M.B., P.C., and R.R. performed experiments and edited the manuscript; M.E.T. provided samples and edited the manuscript; R.N.C designed the study and wrote the manuscript.

---

**Fig. S1.**
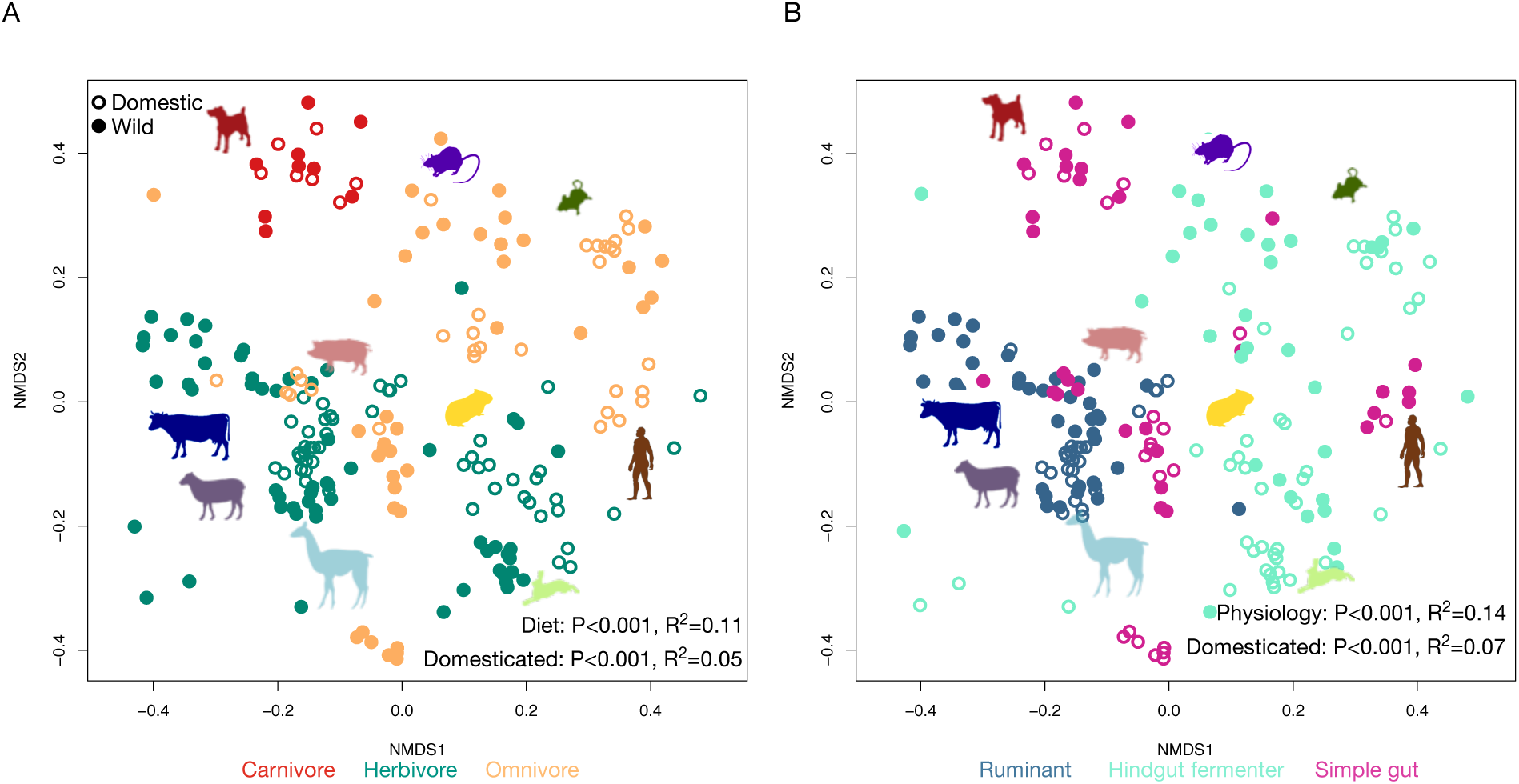
Diet type (**A**) and digestive physiology (**B**) were associated with variation in gut microbial community composition amongst wild (closed circles) and domestic (open circles) mammals, visualized here with nonmetric multidimensional scaling of Bray-Curtis dissimilarity.

**Fig. S2.**
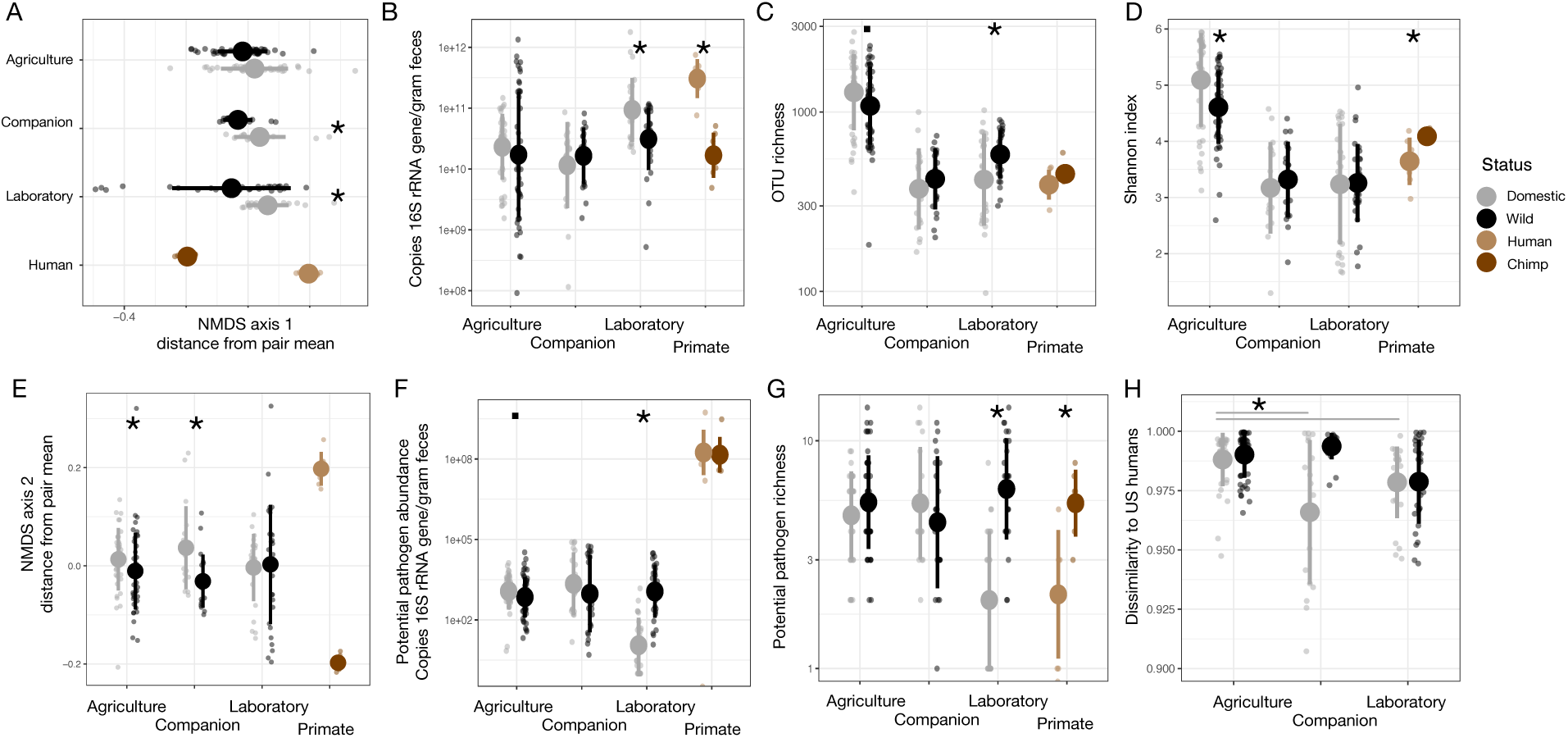
Ordination axis shifts (**A, E**), microbial load (**B**), OTU richness (**C**), Shannon index (**D**), potential pathogen abundance (**F**), and potential pathogen richness (**G**) varied by domestication status for at least one domestication type (agriculture, companion, or laboratory) in cross-species dataset. Trends often mirrored those seen in comparing humans to chimpanzees. (**H**) Bray-Curtis dissimilarity to industrialized humans varied by domestication status and domestication type. Asterisks indicate P<0.05 and periods indicate P<0.1 Mann-Whitney U test.

**Fig. S3.**
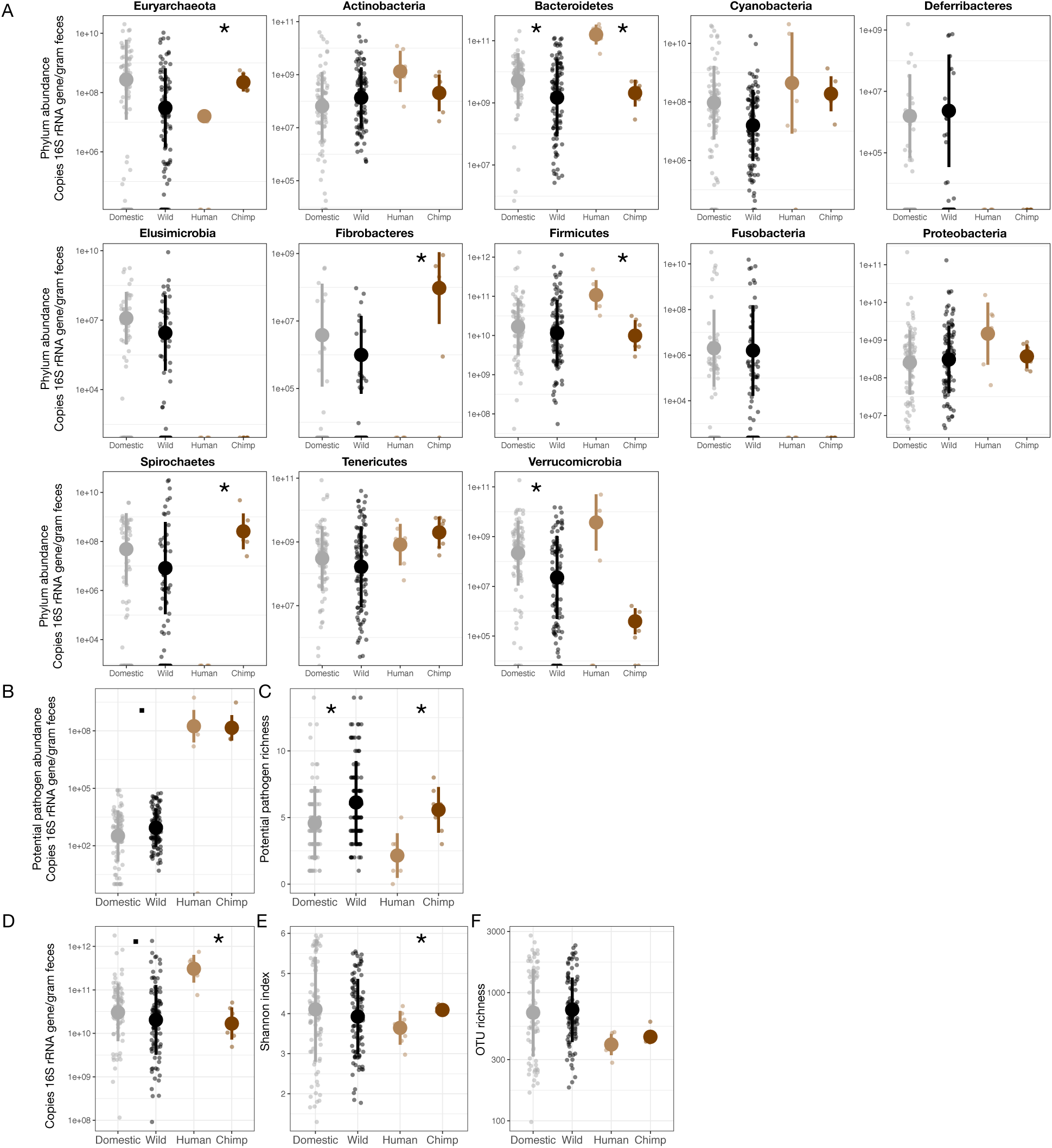
Some phylum abundances (**A**) and potential pathogen community characteristics (**B, C**) varied with domestication in our cross-species dataset. Microbial density (quantified as 16S rRNA gene copies per gram feces; **D**) and alpha-diversity metrics (Shannon index (**E**) and OTU richness (**F**)) did not vary consistently between wild and domestic animals. Asterisks indicate P<0.05 and periods indicate P<0.1 Mann-Whitney U test. Analyses by phylum included Bonferroni multiple hypothesis correction.

**Fig. S4.**
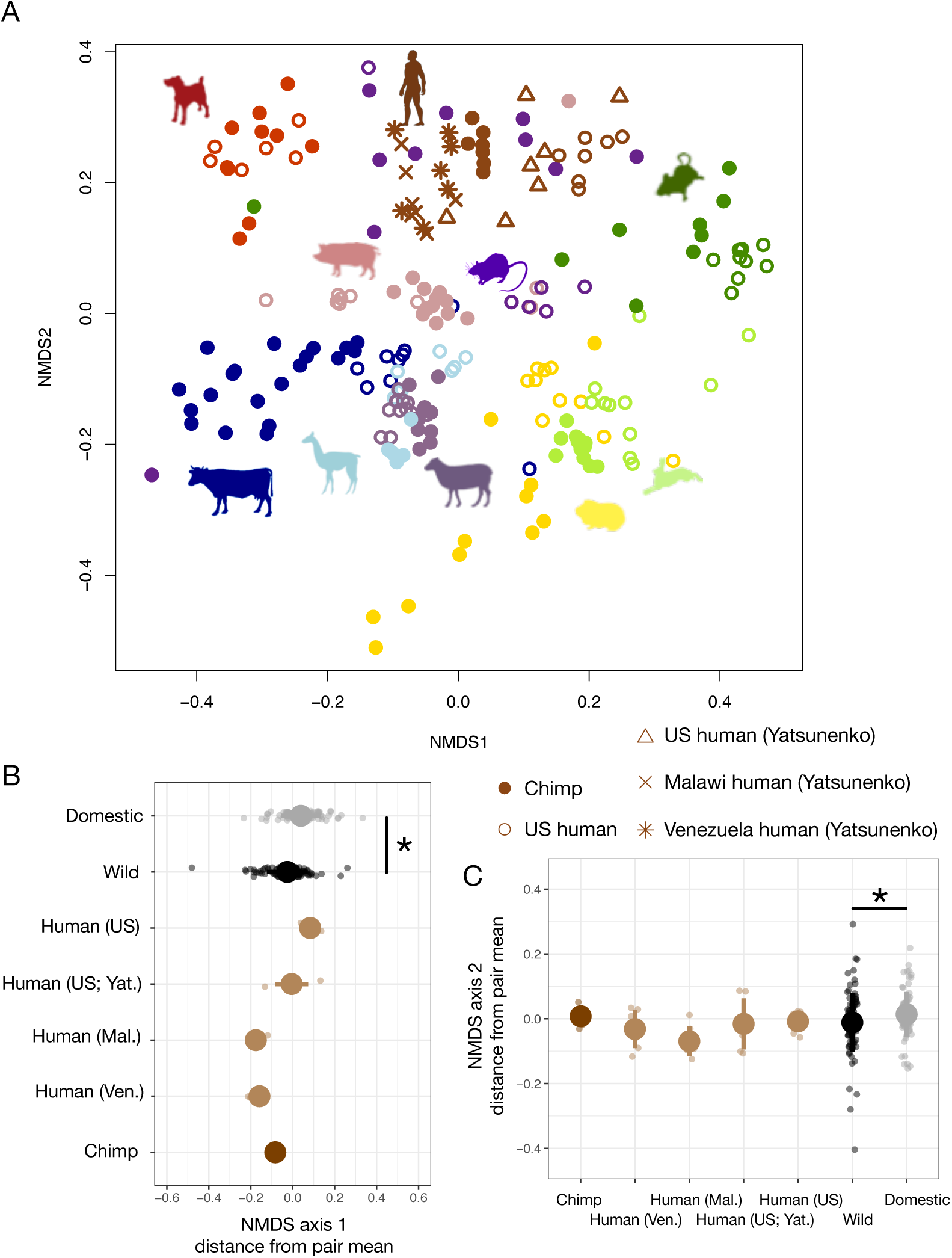
Inclusion of additional human gut microbiota samples shows that while humans and chimpanzees cluster relative to other animals (**A**), traditional human populations do not demonstrate the same shifts along nonmetric multidimensional scaling (NMDS) axis 1 (**B**) and 2 (**C**) as chimpanzees relative to industrialized humans or wild animals relative to domestic animals. NMDS calculated with Bray-Curtis dissimilarity. Asterisks indicate P<0.05 Mann-Whitney U test.

**Fig. S5.**
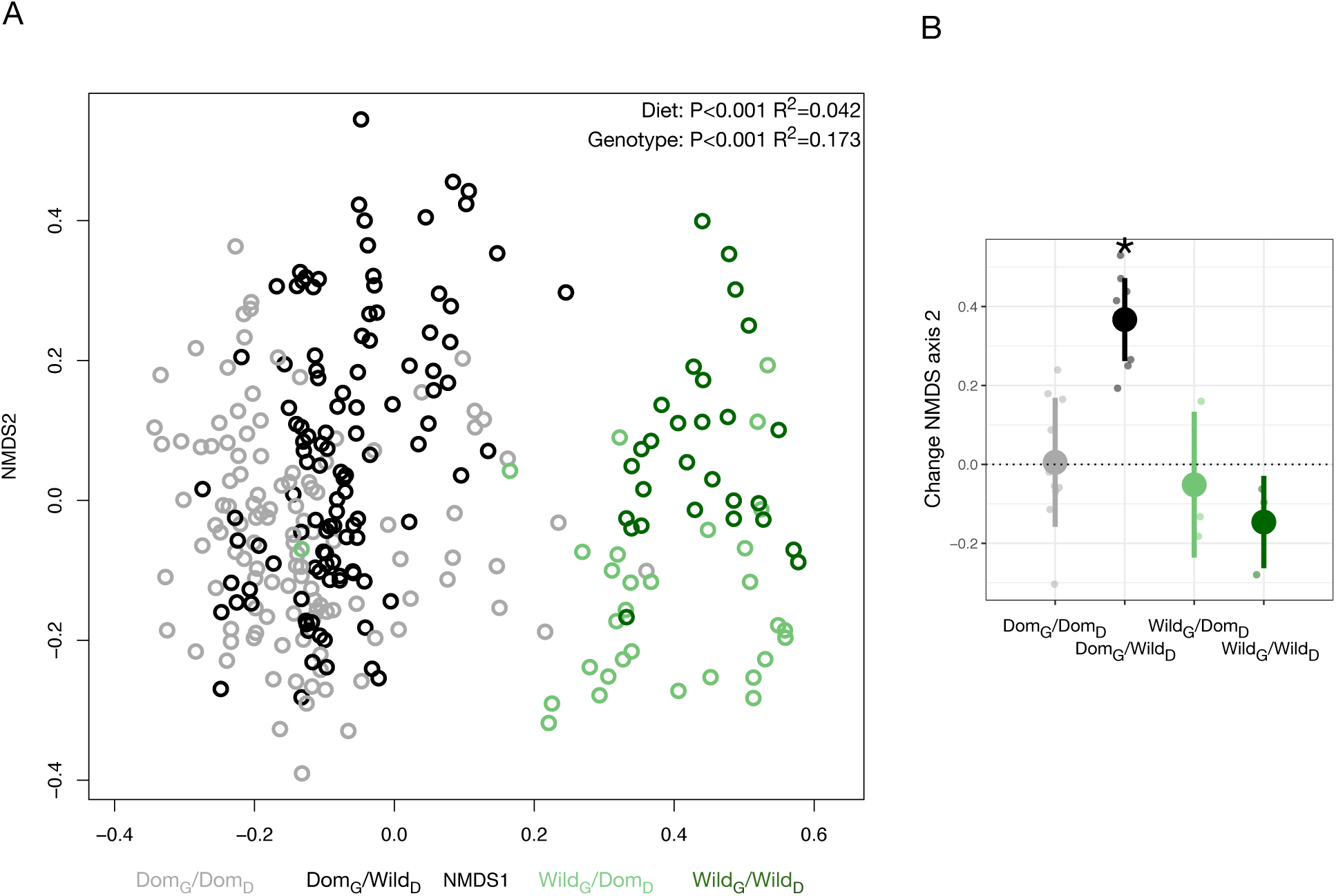
(**A**) Nonmetric multidimensional scaling (NMDS) of all time points illustrates significant effects of genotype and diet on Bray-Curtis dissimilarity. (**B**) Dom_G_/Wild_D_ mice move significantly up along the second NMDS axis between day 0 and 28 of the experiment. Asterisk indicates P<0.05 one-sample Wilcoxon test.

**Fig. S6.**
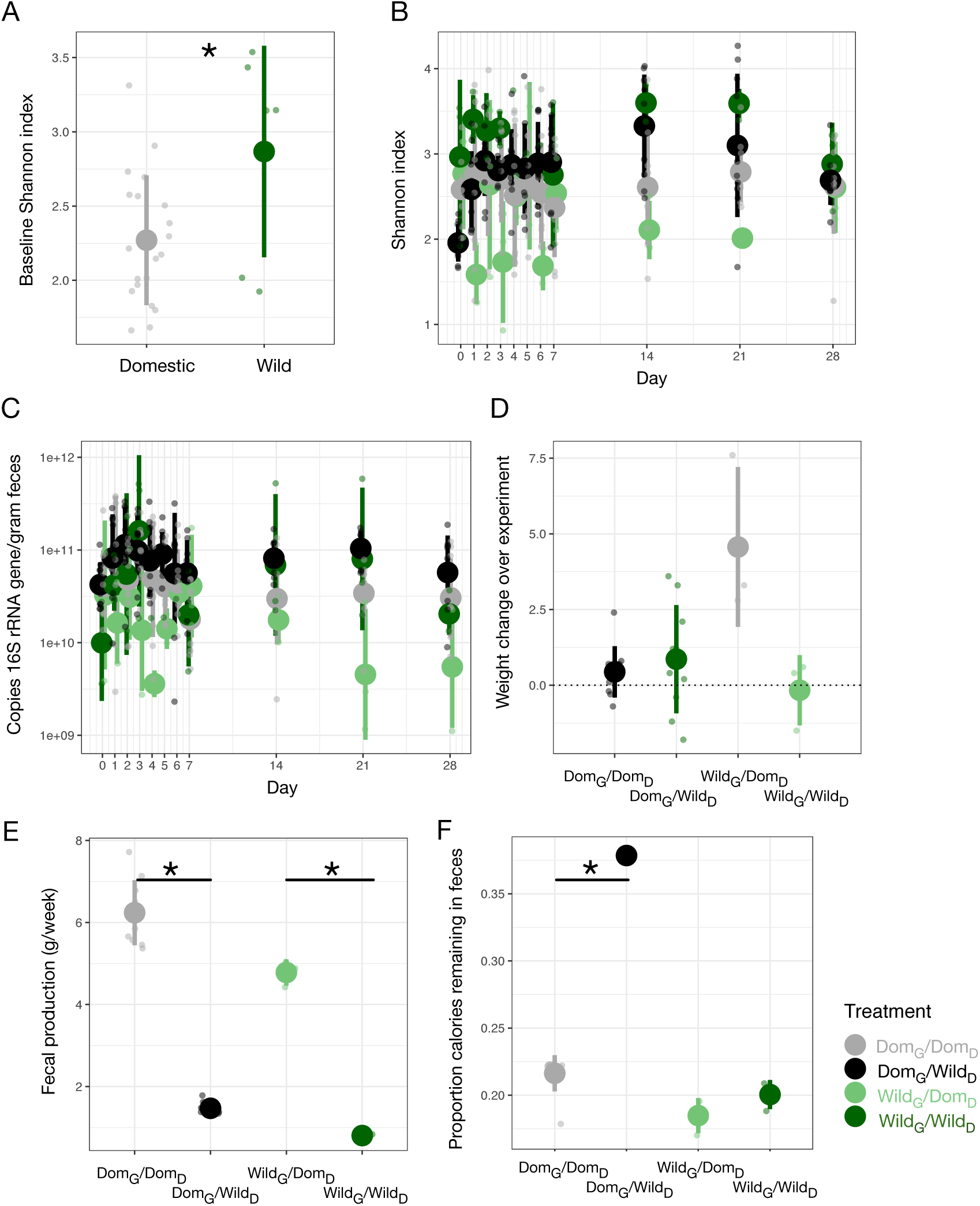
(**A**) Shannon index differed between genotypes on day 0. (**B**) Shannon index plotted by experimental groups over time. (**C**) Microbial load (quantified as 16S rRNA gene copies per gram feces) plotted by experimental groups over time. (**D**) Individual weight gain over the course of the experiment was highest in Wild_G_/Dom_D_ mice. (**E**) Total fecal production over one week differed between experimental groups. (**F**) Calories remaining in feces as a function of total calories consumed varied by diet in Dom_G_ mice. Asterisks in (A, E, F) indicate P<0.05 Mann-Whitney U test.

**Fig. S7.**
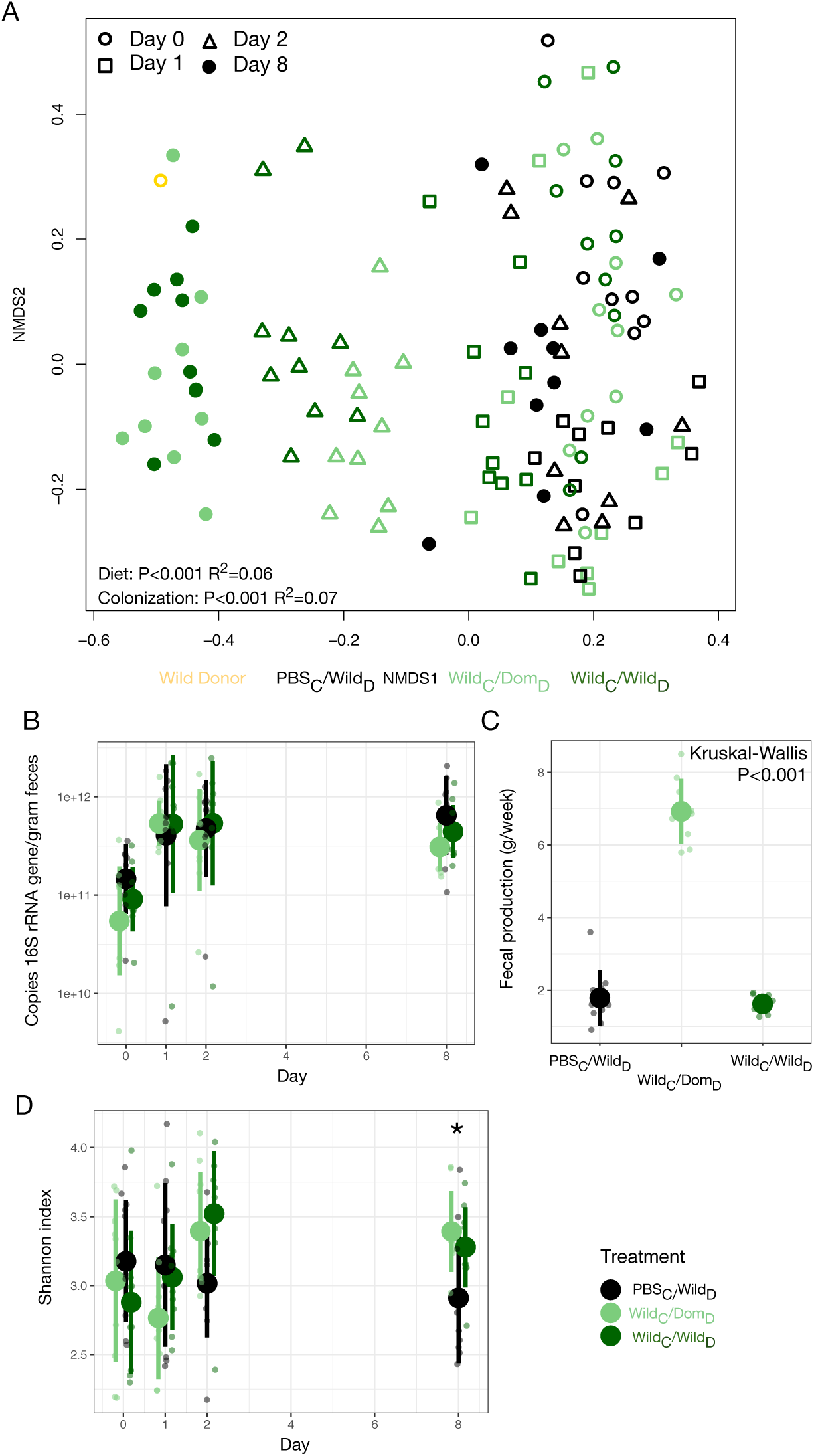
(**A**) Nonmetric multidimensional scaling (NMDS) of all time points illustrated significant effects of colonization and diet treatment on Bray-Curtis dissimilarity. (**B**) Microbial load by experimental groups plotted over time. (**C**) Total fecal production over one week differed between experimental groups. (**D**) Shannon index plotted by experimental groups over time. Asterisks indicate P<0.05 Kruskal-Wallis test.

**Fig. S8.**
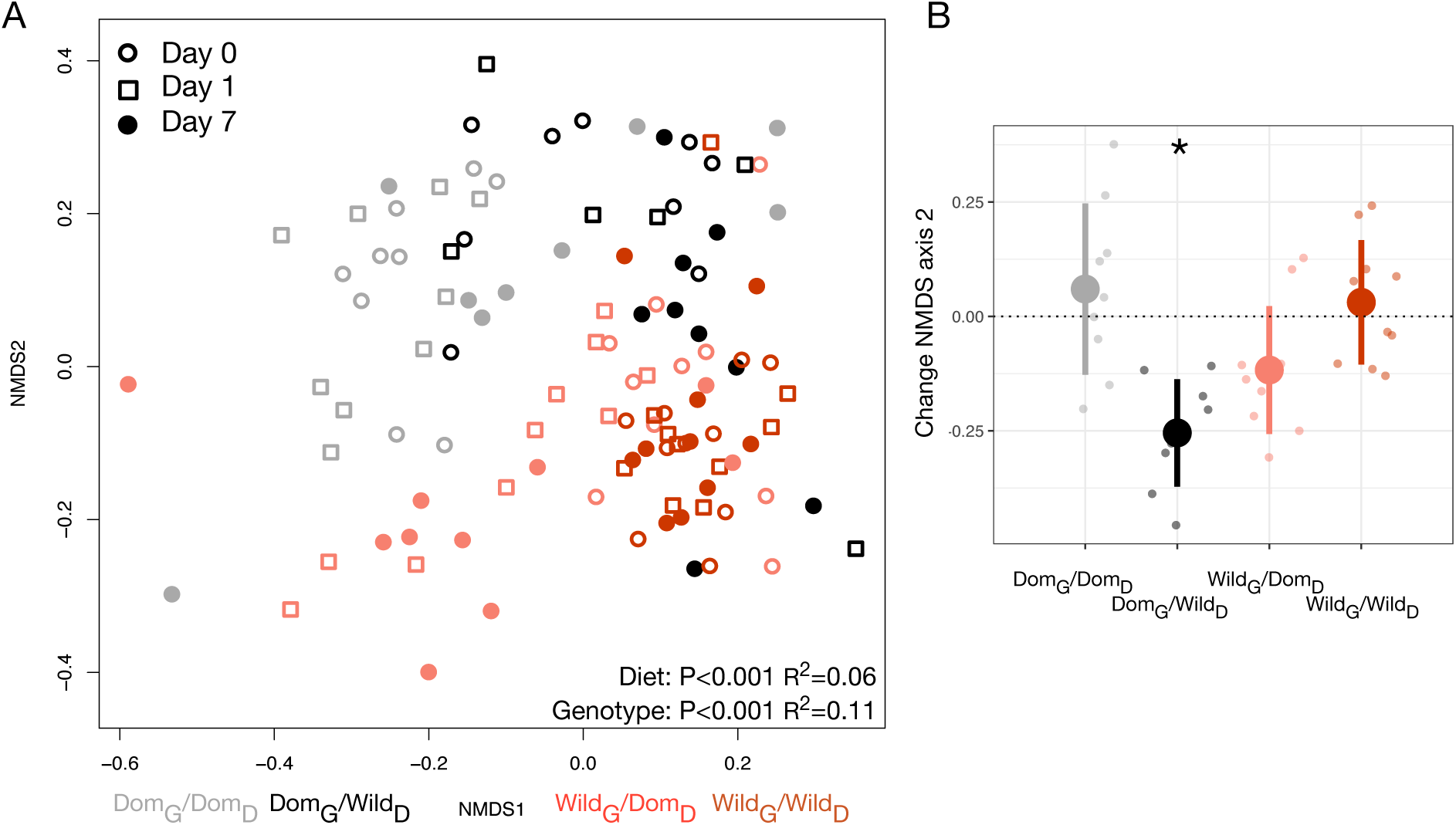
(**A**) Nonmetric multidimensional scaling (NMDS) of all time points illustrated significant effects of genotype and diet on Bray-Curtis dissimilarity. (**B**) Dom_G_/Wild_D_ canids moved significantly down along the second NMDS axis between day 0 and 7 of the experiment. Asterisk indicates P<0.05 one-sample Wilcoxon test.

**Fig. S9.**
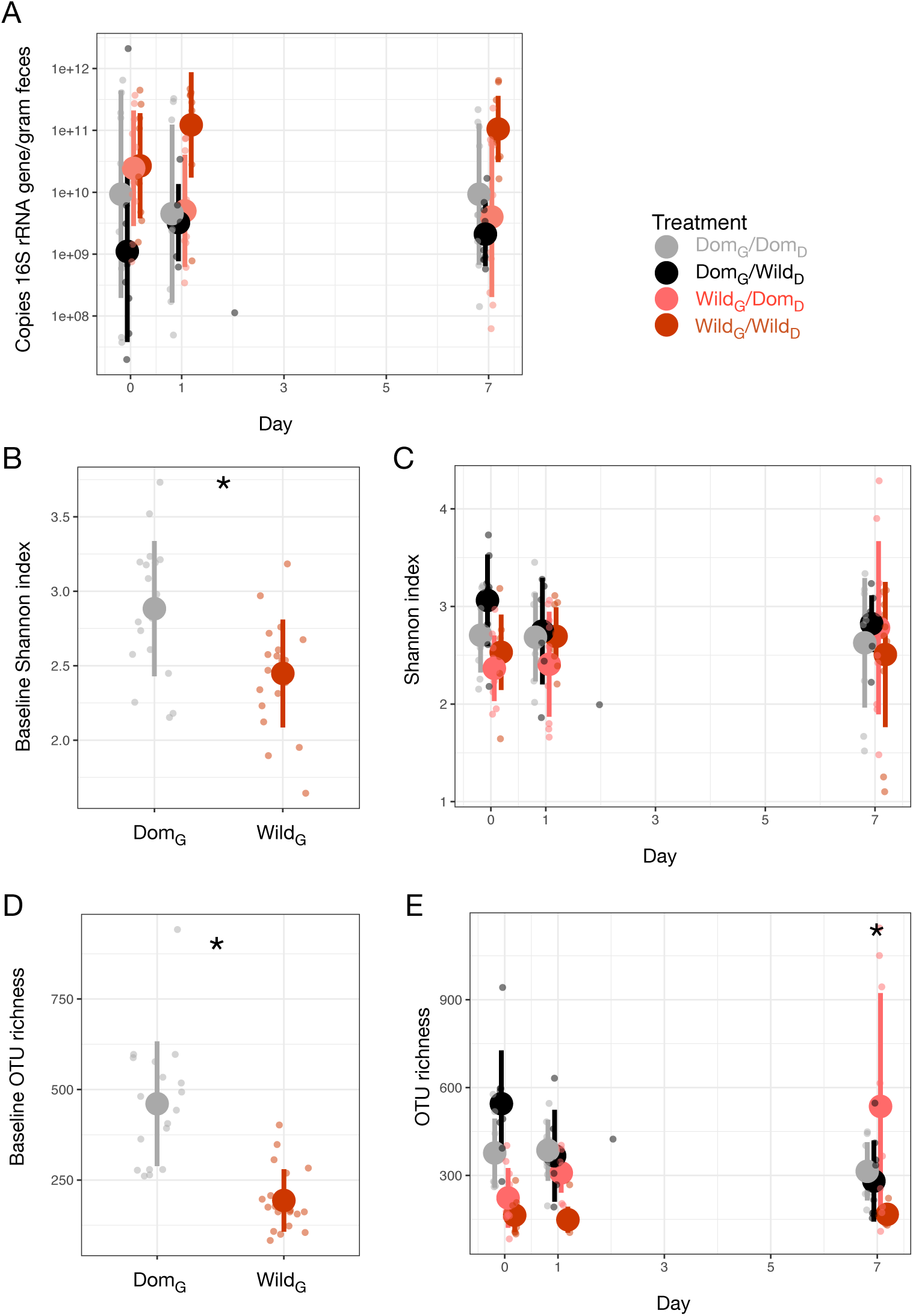
(**A**) Microbial load plotted by experimental groups over time. (**B**) Shannon index differed between genotypes on day 0. (**C**) Shannon index plotted by experimental groups over time. (**D**) OTU richness differed between genotypes on day 0. (**E**) OTU richness plotted by experimental groups over time. Asterisks for (B, D) indicate P<0.05 Mann-Whitney U test. Asterisk for (E) indicates P<0.05 Kruskal-Wallis test.

**Table S1.** Nutritional information for experimental diets.

|  | Mouse |  |  |  | Canid |  |
| --- | --- | --- | --- | --- | --- | --- |
|  | Wild diet |  |  | Domestic diet | Wild diet | Domestic diet |
|  | Mealworms<br>(5% by<br>weight) | Birdseed<br>(95%) | Mix | Prolab Isopro<br>RMH 3000 | Raw chicken | Nutrisource Lamb Meal<br>and Peas Grain Free<br>Diet |
| Crude Protein<br>(min) | 50.0 | 9.0 | 11.1 | 22.0 | 14.1 | 28.7 |
| Crude Fat (min) | 24.0 | 4.0 | 5.0 | 5.0 | 28.7 | 18.3 |
| Crude Fiber (max) | 9.5 | 15.0 | 14.7 | 5.0 | 0.0 | 3.7 |
| Phosphorus (min) | 0.5 |  |  | 0.8 | 0.8 | 1.2 |
| Moisture (max) | 10.0 | 13.0 | 12.9 | 10.0 |  | 9.5 |
| Calories/gram |  |  | 5143 | 4136 | 3190 | 4088 |

## References and Notes

1. C. De Filippo et al., Impact of diet in shaping gut microbiota revealed by a comparative study in children from Europe and rural Africa. PNAS 107, 14691–14696 (2010).

2. A. H. Moeller et al., Rapid changes in the gut microbiome during human evolution. PNAS 111, 16431–16435 (2014).

3. A. H. Moeller, The shrinking human gut microbiome. Curr Opin Microbiol 38, 30–35 (2017).

4. S. A. Smits et al., Seasonal cycling in the gut microbiome of the Hadza hunter-gatherers of Tanzania. Science 357, 802–806 (2017).

5. R. E. Ley, P. J. Turnbaugh, S. Klein, J. I. Gordon, Microbial ecology: human gut microbes associated with obesity. Nature 444, 1022–1023 (2006).

6. L. M. Cox et al., Altering the intestinal microbiota during a critical developmental window has lasting metabolic consequences. Cell 158, 705–721 (2014).

7. N. Kamada, S. U. Seo, G. Y. Chen, G. Nunez, Role of the gut microbiota in immunity and inflammatory disease. Nature Rev Immunol 13, 321–335 (2013).

8. M. A. Zeder, The domestication of animals. J Anthropol Res 68, 161–190 (2012).

9. C. Theofanopoulou et al., Self-domestication in Homo sapiens: Insights from comparative genomics. PLoS ONE 12, e0185306 (2017).

10. L. A. David et al., Diet rapidly and reproducibly alters the human gut microbiome. Nature 505, 559–563 (2014).

11. R. N. Carmody et al., Diet dominates host genotype in shaping the murine gut microbiota. Cell Host Microbe 17, 72–84 (2015).

12. J. M. Allen et al., Exercise alters gut microbiota composition and function in lean and obese humans. Med Sci Sports Exer 50, 747–757 (2018).

13. E. V. Lamoureux, S. A. Grandy, M. G. I. Langille, Moderate exercise has limited but distinguishable effects on the mouse microbiome. mSystems 2, e00006–00017 (2017).

14. K. A. Dill-McFarland et al., Close social relationships correlate with human gut microbiota composition. Sci Rep 9, 703 (2019).

15. R. E. Antwis, J. M. D. Lea, B. Unwin, S. Shultz, Gut microbiome composition is associated with spatial structuring and social interactions in semi-feral Welsh Mountain ponies. Microbiome 6, 207 (2018).

16. N. A. Bokulich et al., Antibiotics, birth mode, and diet shape microbiome maturation during early life. Sci Trans Med 8, 343ra382 (2016).

17. I. Cho et al., Antibiotics in early life alter the murine colonic microbiome and adiposity. Nature 488, 621–626 (2012).

18. C. Li et al., Effect of early weaning on the intestinal microbiota and expression of genes related to barrier function in lambs. Front Microbiol 9, 1431 (2018).

19. A. S. Wilkins, R. W. Wrangham, W. T. Fitch, The “domestication syndrome” in mammals: a unified explanation based on neural crest cell behavior and genetics. Genetics 197, 795–808 (2014).

20. B. D. Muegge et al., Diet drives convergence in gut microbiome functions across mammalian phylogeny and within humans. Science 332, 970–974 (2011).

21. S. P. Rosshart et al., Wild mouse gut microbiota promotes host fitness and improves disease resistance. Cell 171, 1015–1028 (2017).

22. L. K. Beura et al., Normalizing the environment recapitulates adult human immune traits in laboratory mice. Nature 532, 512–516 (2016).

23. T. Yatsunenko et al., Human gut microbiome viewed across age and geography. Nature 486, 222–227 (2012).

24. S. J. Song et al., Cohabiting family members share microbiota with one another and with their dogs. eLife 2, e00458 (2013).

25. R. N. Carmody, in Chimpanzees & Human Evolution, M. N. Muller, R. W. Wrangham, D. R. Pilbeam, Eds. (Harvard University Press, Cambridge, MA, 2017), pp. 311–338.

26. E. D. Sonnenburg et al., Diet-induced extinctions in the gut microbiota compound over generations. Nature 529, 212–215 (2016).

27. E. Axelsson et al., The genomic signature of dog domestication reveals adaptation to a starch-rich diet. Nature 495, 360–364 (2013).

28. K. D. Kohl, R. B. Weiss, J. Cox, C. Dale, M. Denise Dearing, Gut microbes of mammalian herbivores facilitate intake of plant toxins. Ecol Lett 17, 1238–1246 (2014).

29. M. G. Dominguez Bello, R. Knight, J. A. Gilbert, M. J. Blaser, Preserving microbial diversity. Science 362, 33–34 (2018).

30. S. M. Hird, Evolutionary biology needs wild microbiomes. Front Microbiol 8, 725 (2017).

31. A. Alberdi, O. Aizpurua, K. Bohmann, M. L. Zepeda-Mendoza, M. T. P. Gilbert, Do vertebrate gut metagenomes confer rapid ecological adaptation? Trends Ecol Evol 31, 689–699 (2016).

32. M. Combs, E. E. Puckett, J. Richardson, D. Mims, J. Munshi-South, Spatial population genomics of the brown rat (Rattus norvegicus) in New York City. Mol Ecol 27, 83–98 (2018).

33. J. S. Weese, T. Shury, M. D. Jelinski, The fecal microbiota of semi-free-ranging wood bison (Bison bison athabascae). BMC Vet Res 10, 120 (2014).

34. E. Schwarz, H. K. Schwarz, The wild and commensal stocks of the house mouse, Mus musculus Linnaeus. J Mammal 24, 59–72 (1943).

35. J. G. Caporaso et al., Global patterns of 16S rRNA diversity at a depth of millions of sequences per sample. PNAS 108, 4516–4522 (2011).

36. J. G. Caporaso et al., Ultra-high-throughput microbial community analysis on the Illumina HiSeq and MiSeq platforms. ISME J 6, 1621–1624 (2012).

37. C. F. Maurice, H. J. Haiser, P. J. Turnbaugh, Xenobiotics shape the physiology and gene expression of the active human gut microbiome. Cell 152, 39–50 (2013).

38. J. Oksanen et al. vegan: Community Ecology Package. R package version 2.4-2. https://CRAN.R-project.org/package=vegan (2017).

39. S. W. Kembel et al., Architectural design influences the diversity and structure of the built environment microbiome. ISME J 6, 1469–1479 (2012).

40. A. T. Reese et al., Urban stress is associated with variation in microbial species composition—but not richness—in Manhattan. ISME J 10, 751–760 (2016).

